# Predictive Metabolic Pathways of Lactic Acid Bacterial Strains Isolated from Fermented Foods

**DOI:** 10.1101/2020.07.01.181941

**Authors:** Pynhunlang Kharnaior, Prakash M. Halami, Jyoti Prakash Tamang

**Affiliations:** DAICENTRE (DBT-AIST International Centre for Translational and Environmental Research) and Bioinformatics Centre, Department of Microbiology, School of Life Sciences, Sikkim University, Gangtok 737102, Sikkim, India; CSIR-Central Food Technological Research Institute, Microbiology and Fermentation Technology, Mysuru, 570020, Karnataka, India

**Author notes:** Corresponding author: Professor Jyoti Prakash Tamang.

**Keywords:** Predictive functionality, Metabolic Pathways, Fermented Foods, LAB, PICTRUSt

## Abstract

We attempted to use PICTRUSt2 software and bioinformatics tool to infer the raw sequences obtained from pure strains of *Lactococcus lactis* and *Lactobacillus plantarum* isolated from some fermented foods in India, which were identified by 16S rRNA gene sequencing method. Predictive metabolic pathways of 16S sequences of LAB strains were predicted by PICRUSt2 mapped against KEGG database, which showed genes associated with metabolism (36.74%), environmental information processing (32.34%), genetic information processing (9.86%) and the unclassified (21.06%). KGGE database also showed the dominant genes related to predictive sub-pathways of metabolism at level-2 were membrane transport (31.16%) and carbohydrate metabolism (12.42%).

## Introduction

Ethnic fermented foods of India have socio-cultural values, rich gastronomy and also contribute health benefits to consumers (Tamang 2020), and are composed by diverse types of bacteria, fungi and yeasts (Tamang et al. 2012). Lactic acid bacteria present in fermented foods have several functional properties, therapeutic uses and health promoting benefits (Tamang et al. 2016; Goel et al. 2020). Phylogenetic investigation of communities by reconstruction of unobserved states (PICRUSt2), an omics-based machine learning software in bioinformatics (Douglas et al. 2019), is commonly used to predict the genes markers of sequences generated by high-through sequences and shot-gun sequences from food samples (Tamang et al. 2020). However, there is a limited application of PICTRUSt to predict functional metabolic pathways in sequences obtained from isolated bacterial strains (Medvecky et al. 2018). In the present study, we analysed sequence of five lactic acid bacteria isolated from Indian fermented and sun-dried foods to predict the metabolic pathways by bioinformatics tools.

## Materials and Methods

Three different types of Indian fermented foods were collected from Sikkim and Karnataka states in India viz. *sidra* (dried fish products of Sikkim), *kinema* (fermented soybean food of Sikkim), and *dahi* (fermented milk product of Karnataka and Sikkim). Ten gram each of samples was homogenized in 90 mL of 0.85% physiological saline using stomacher lab blender 40 (Seward, United Kingdom) for few min, and were plated MRS (Man-Rogosa-Sharpe) agar (M641, HiMedia, Mumbai, India), and incubated at 30°C for 24–48 h. About 60 isolates were randomly isolated and preliminarily tested for Gram-stain, catalase test, arginine hydrolysis and other phenotypic characteristics (Pradhan and Tamang 2019). Out of 60 isolates, five isolates were grouped on the basis of phenotypic characteristics viz. isolate FS2 *(sidra),* C2D (*dahi*-Mysore), DHCU70 *(dahi,* Gangtok), SP2C4 *(kinema)* and KP1 *(kinema).*

Genomic DNA of 5 strains were extracted following the method of Cheng and Jiang (2006). Amplified 16S rDNA was obtained from each strain by polymerase chain reaction (PCR) with the universal primers in a Thermal cycler (Applied Biosystems-2720, USA). Sequencing of the amplicons was performed in an automated DNA Analyzer (ABI 3730XL Capillary Sequencers, Applied Biosystems, Foster City, CA, USA). Taxonomy of bacterial strains was assigned by comparing the DNA sequences in NCBI (National Center for Biotechnology Information) database using BLAST (basic local alignment search tool) 2.0 program and the phylogenetic tree was constructed by neighbor-joining method using MEGA7.0 software (Kumar et al. 2016).

## Results and Discussion

The 16S rRNA gene sequences of five strains were analysed for the functionality using PICRUSt2-algorithm (Douglas et al. 2019). The sequences were aligned for placement of the study sequences into a reference phylogeny, and the gene copy number of test sequences were effectuated at default parameters (Barbera et al. 2019). Annotation of gene function and pathways inference with high level function (Ye and Doak 2009) were mapped against KEGG (Kyoto Encyclopaedia of Genes and Genomes) database (Kanehisa et al. 2017). Raw reads were normalized into relative abundances and data visualization using MS-Excel 365.

Based on results of 16S rRNA gene sequences, a phylogenetic tree was constructed using neighbour-joining for identification of bacterial strains (Fig. 1). FS2 strain isolated from *sidra,* C2D strain isolated from *dahi* and SP2C4 isolated from *kinema* were identified as *Lactococcus lactis,* and DHCU70 strain isolated from *dahi* and KP1 strain isolated from *kinema* were identified as *Lactobacillus plantarum* (Fig. 1). The functional features of sequences of 5 strains of LAB obtained from 16S rRNA data were predicted by using PICRUSt2 mapped against KEGG database. The overall functional features of LAB strains showed genes associated with metabolism (36.74%), environmental information processing (32.34%), genetic information processing (9.86%) and the unclassified (21.06%) at Level-1 (Fig. 2a). At the Level-2, Dominant genes related to predictive sub-pathways of metabolism at level-2 were membrane transport (31.16%) and carbohydrate metabolism (12.42%) (Fig. 2b). KGGE database showed tyrosine metabolism was dominant in *Lactococcus lactis* (Fig. 3a), indicating its role in flavour development in the product since tyrosine is an aromatic amino acids (Parthasarathy et al., 2018). Predictive genes in *Lb*. *plantarum* showed the folate biosynthesis (Fig. 3b), that can possibly play an important key in the development of drug against infectious disease (Bertacine Dias et al. 2018). Phosphotransferases system (PTS), the source of transport and phosphorylation of various sugar and other sugar derivatives in bacteria (Deutscher et al. 2006), was detected only in *Lb*. *plantarum.* We did not detect any predictive genes for human diseases in *Lb. plantarum* and *Lc. lactis* strains. The ATP-binding cassette transporters (ABC transporters) genes were dominant in all five strains, which facilitate the selective uptake of di- and tripeptides (Doeven et al. 2005). We can conclude that predictive metabolic pathways (Baranwal et al. 2020) can be inferred more in details of even isolated strains by KGGE database which may help to understand functionality of the LAB strains in foods.

**Fig. 1:**
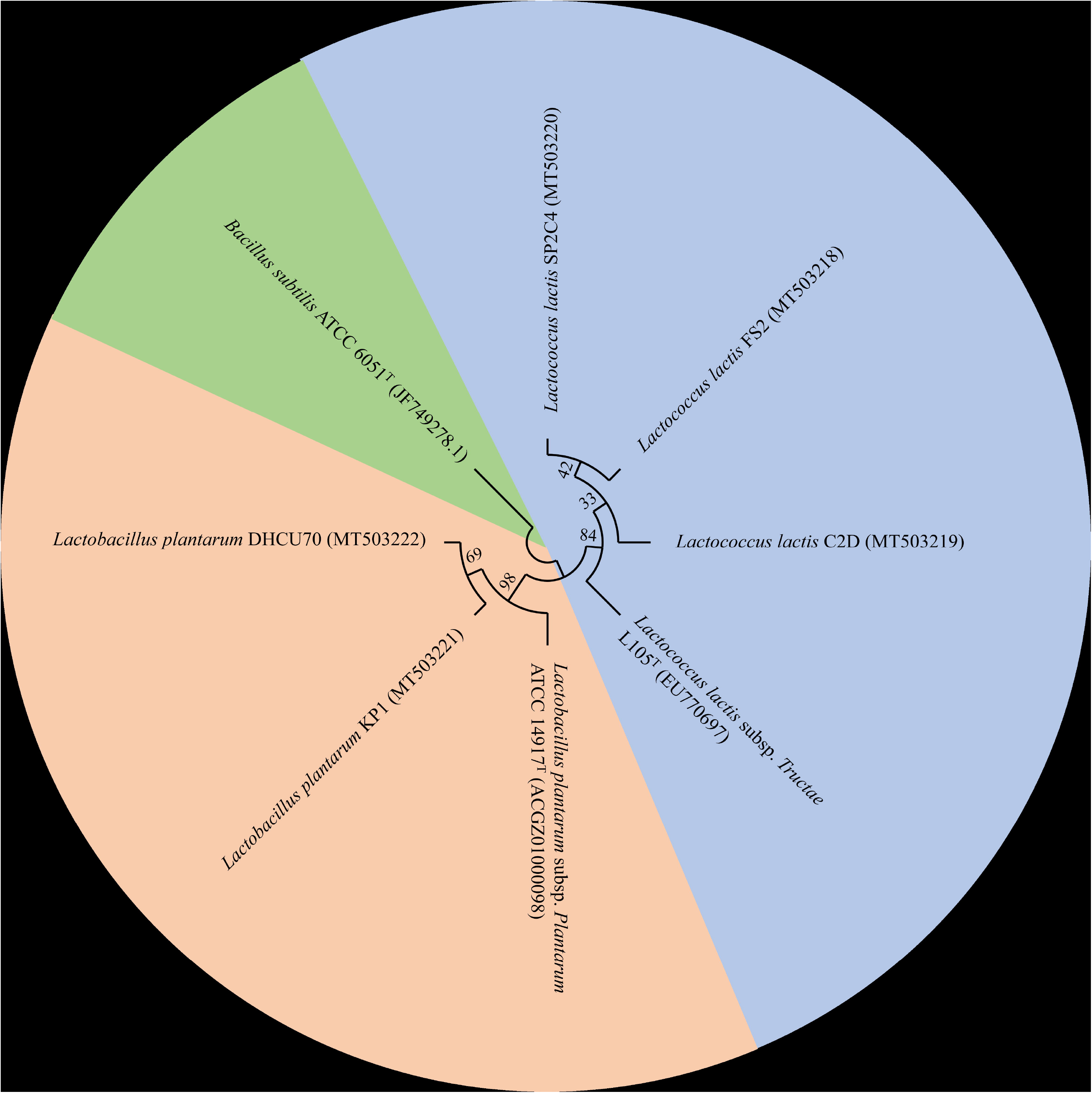
Neighbor-joining phylogenetic analysis using MEGA7 based on 16S rRNA gene sequences of lactic acid bacterial strains (FS2, C2D, SP2C4, KP1 and DHCU70) isolated from fermented foods with bootstrap values (expressed as percentage of 1000 replicates). *Bacillus subtilis* ATCC 6051T (Genbank accession number JF749278.1) was used as an outgroup.

**Fig. 2:**
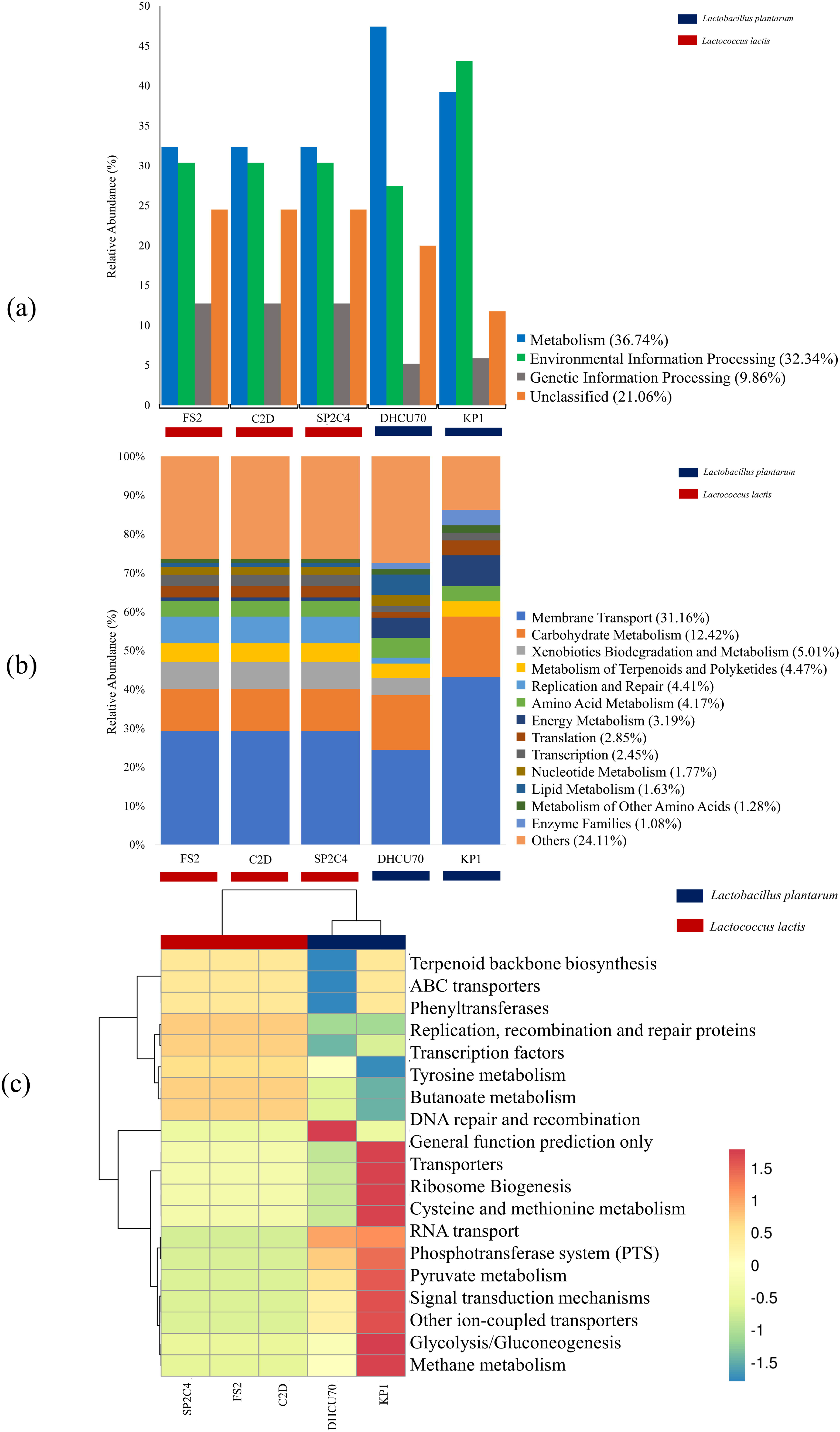
Overall predictive functional features of LAB strains via PICRUSt2 mapped against KEGG database. A bar-plot for (a) Level-1 and (b) Level-2, and a heatmap of predominant pathways (c) Level-3 were plotted.

**Fig. 3:**
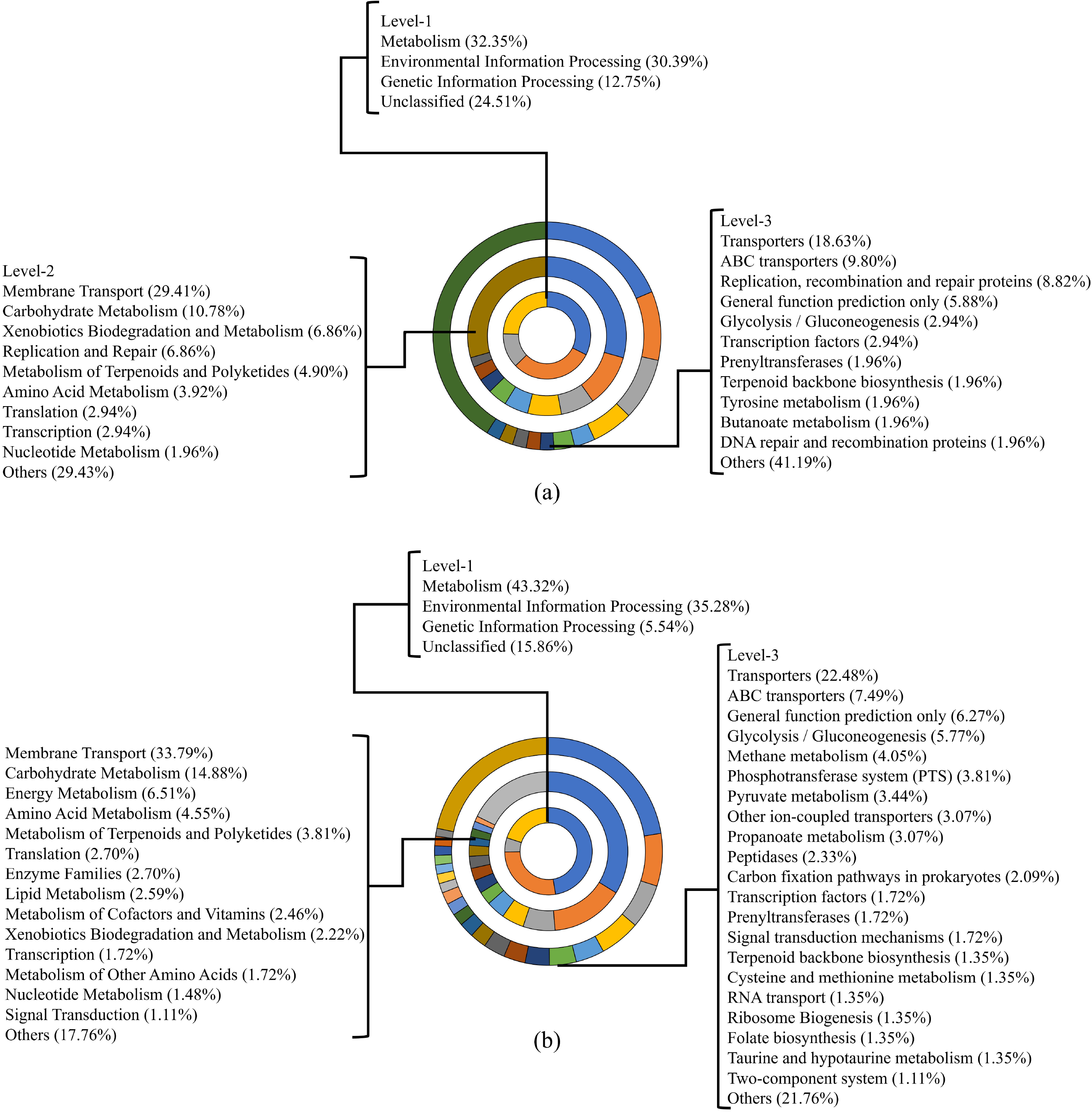
Doughnut representation were plotted in comparison between the two species identified (a) *Lactococcus lactis* and (b) *Lactobacillus plantarum.*

## Acknoledgement

This work was supported by Department of Biotechnology, Government of India for Twinning Research, project no. BT/PR16706/NER/95/259/2015.

## COMPLIANCE WITH ETHICAL STANDARDS

### Statement on the welfare of animals

This article does not contain any studies involving animals or human subjects performed by any of the authors.

## DATA AVAILABILITY

The sequences retrieved from the 16S rRNA sequencing were deposited at GenBank-NCBI under the nucleotide accession number: MT503218, MT503219, MT503220, MT503221 and MT503222.

